# Robin Hood: non-fitting, non-smoothing image detrending for bleaching correction

**DOI:** 10.1101/667824

**Authors:** Rory Nolan, Maro Iliopoulou, Christian Siebold, E. Yvonne Jones, Sergi Padilla-Parra

**Affiliations:** Cellular Imaging Group, Wellcome Centre for Human Genetics, University of Oxford, Oxford, UK; Division of Structural Biology, University of Oxford, Wellcome Centre for Human Genetics, Headington, Oxford OX3 7BN, UK; Dynamic Structural Virology Group, Biocruces Health Research Institute, 48903 Barakaldo, Spain; Ikerbasque, Basque Foundation for Science, Bilbao, Spain

## Abstract

Recent advances in protein labelling—gene tagging with CRIPSR-Cas9—have made it possible to label proteins of interest endogenously. This represents a major breakthrough in the field of quantitative microscopy, especially when quantifying protein-protein interactions. This is because over-expression of labelled proteins may cause a distortion in localization, function and perhaps artificially force protein-protein interactions due to crowding effects. A microscopy technique that is particularly well suited to detect protein interactions with low photon budgets is number and brightness (N&B). Detrending (removal of global trends in data) is a necessary pre-processing step to N&B calculations, but all current detrending methods perform poorly at low intensities. Here, we present the *Robin Hood* automatic detrending algorithm which performs well at low intensities, evaluating it with simulated and low photon budget live cell images. RH is available as an ImageJ plugin and as an R package.

**STATEMENT OF SIGNIFICANCE:** Fluorescence microscopy in general and Fluorescence fluctuation methods in particular are very much dependent on detrend algorithms and so far, the user needs to decide an arbitrary number to correct for bleaching when using the box plot or the running average approach. Here, we have developed a tool available to everybody as an ImageJ plugin that is automatic, user-free and able to correct bleached images with very low photon counts.

## INTRODUCTION

Many calculations in the fields of fluorescence fluctuation and correlation spectroscopy (FFS^1^ and FCS^2^) rely on the assumption of constant mean intensity of the illuminated sample. Due to many factors possibly including but not limited to photobleaching, sample (cell) movement (both in *xy* and in *z*), laser power fluctuations and detector sensitivity fluctuations, this assumption is invalid. The purpose of *detrending* algorithms is to correct image series (videos) such that the mean intensity of the sample is approximately constant (i.e. without any global *trend*).

Thus far, all detrending methods have been based on fitting^3^ or smoothing^4^. Both of these approaches have serious caveats. Fitting assumes that the fluorescence intensity decay has a certain form. Unpredictable factors (such as those listed above) mean that no particular decay form can be assumed (see example of real biological data in figure 2 which cannot be well-fitted with any conventional function). Moreover, both fitting and smoothing fail when the data cannot be approximated as continuous (fitted and smoothed lines are continuous approximations of data). This is a limitation when dealing with very dim samples and live cells. Fluorescence intensity data at low intensities – where most pixel values are either 0 or 1 – are quasi-binary and hence a continuous approximation does not make sense (see figure 2 for an example of fitting a line to binary data). This means that neither fitting nor smoothing are applicable detrending methods at low intensities. This is the crucial caveat of these methods because, when bleaching is a problem, it is common to reduce laser power to reduce bleaching, which leads directly to lower intensity images. With fitting and smoothing techniques, it may sometimes be advisable to increase the laser power to achieve higher intensities such that the detrending routines will function properly. This means one may need to bleach more in order to be able to properly correct for bleaching. This farcical situation further warrants a new detrending technique which can function at low intensities. Here we present the *Robin Hood* (RH) detrending algorithm which does not rely on fitting or smoothing. The basic idea is to *take* intensity counts from frames of greater than average intensity and *give* them directly to frames of less than average intensity. This has the desired effect of detrending the image series because both the giver and receiver frames of each count *swap* end up closer to the mean intensity than they were before the swap took place.

## METHODS

### Robin Hood Algorithm

The method for deciding which swaps to make is as follows:

1. Calculate the mean intensity *μ* of the entire image series.
2. For each frame *i*, calculate the mean intensity *μ*_*i*_ of that frame.
3. For each frame *i*, calculate the distance between the mean intensity of that frame and the mean intensity of the image series: *d*_*i*_ = *μ*_*i*_ − *μ*. Frames with *μ*_*i*_ > *μ* will have *d*_*i*_ > 0 and frames with *μ*_*i*_ < *μ* will have *d*_*i*_ < 0.
4. Frames have probability weights *d*_*i*_ to be taken from and probability −*d*_*i*_ to be given to (where negative weights are interpreted as zero). Frames which start with mean frame intensity greater than *μ* may not give away so many counts as to go to having mean intensity less than *μ* and frames which start with mean frame intensity less than *μ* may not get so many counts as to go to having mean intensity greater than *μ* (this limits the total number of swaps that can happen, putting an upper bound on the running time of the algorithm).
5. When a giving frame *i* and a receiving frame *j* are chosen, choose an intensity count at random (each count is equally likely) to take from frame *i*. This count will be at some pixel position *p* in frame *i*. Give that count to pixel position *p* in frame *j* (so counts only travel *along* any given pixel). Mathematically, this amounts to subtracting 1 from pixel position *p* in frame *i* and adding 1 to pixel position *p* in frame *j*.

Point 5 above ensures that the mean intensity projection of the detrended image is equal to that of the original image. This is important because the mean intensity (or summed intensity) projection is commonly used as a summary image for the whole image series, so to conserve it is highly preferable. RH is the only detrending algorithm which guarantees this conservation. With single-point FCS traces, it is impossible to calculate the mean of a frame because there is no *frame*. Hence, there is no way to get a summary statistic for the mean intensity at a given time point, so point 2 above cannot be completed and hence RH cannot be applied to single-point traces. It is, however, perfectly applicable to single-line scanning FCS.

The number of swaps to make is determined with our 2016 algorithm for choosing detrending parameters.^5^ Note that this algorithm assumes that the image is in units of photon counts and as such RH can only be used on such images.

### Mathematical Simulations

We have developed an R package called *brownded* (https://github.com/rorynolan/brownded) for simulating bounded brownian motion in any number of dimensions, where *bounded* brownian motion is brownian motion in an *n*-dimensional box where the particles collide elastically (without loss of energy) with the boundaries of the box. *brownded* allows specification of the number of dimensions, the number of particles, the size of the box and the diffusion coefficient of the particles. *brownded* also facilitates the simulation of images created from fluorescent particles undergoing bounded *brownian* motion. It allows specification of the time at which each image should be taken, the pixel size and the brightness of the particles. Each fluorescent particle contributes photon counts to its pixel of residence at that time according to a poisson process. Finally, *brownded* facilitates the synthetic bleaching of fluorescent particles so bleaching can be investigated with images produced with *brownded*.

### Detrending algorithm evaluation

The general idea for evaluating a given detrending algorithm was:

1. Simulate a number *N* of particles diffusing with known diffusion rate.
2. Simulate photon emission from these particles with chosen brightness *ϵ* and create an image series from this, being careful to (virtually) sample at a rate appropriate for number and brightness analysis.
3. Bleach the simulation with a chosen constant bleaching rate.
4. Simulate photon emission from the bleached simulation (bleached particles don’t emit photons) with the same brightness *ϵ* and create an image series.
5. Detrend the bleached image series.
6. Evaluate the detrending algorithm by measuring how close the brightness of the detrended bleached image series is to the known simulated brightness.

This was done with *N* = 10,000 particles with 21 replicates (I chose this many replicates to limit the simulations to two weeks) of every combination of brightness *ϵ* = 0.001, 0.01, 0.1 and bleaching fractions of 0%, 1%, 5%, 10%, 15%, 20%, 25%, 30%. Images were 64×64 pixels and 5,000 frames. The performance was evaluated using the *mean relative error*. For a given brightness and bleaching fraction, mean relative error = |(calculated brightness after detrending) − (true brightness)|(true brightness)

### Cell Culture

COS-7 cells were cultured in DMEM supplemented with 10% fetal bovine serum, 1% penicillin-streptomycin, and 1% L-glutamine to give DMEM^complete^ medium. All cells were maintained in a 37°C incubator supplied with 5% CO_2_. Cells were imaged in complete DMEM without Phenol red in Ibidi imaging chambers (Gmbh, Germany).

### Reagents and antibodies

CCL5/RANTES was included in Human Chemokine Protein Sampler Pack from R&D systems..

### Plasmids Transfections

hCCR5-mTFP1 is deposited in Addgene/ Padilla-Parra Lab (https://www.addgene.org/110193/) it was transiently transfected in COS-7 cells using GeneJuice according to the manufacturer’s protocol (Novagen). Transiently-transfected cells were analysed by confocal LSM equipped with photon counting detectors 48 hours later.

### Microscopy acquisitions

Live COS-7 cells expressing CCR5-mTFP1 were imaged using a SP8 X SMD confocal microscope Leica microscope from Leica Microsystems (Manheim, Germany). Cells of interest were selected under a 63x oil-immersion objective. CCR5-mTFP1 cells were excited using the White Light Laser (Leica) at 470nm tuned at 80MHz.Subsequently detected by hybrid detectors in photon counting mode. Time-lapse acquisitions of 500 frames in 256 ×256 pixels, while the dwell time was 2.43µs and the frame rate was 1.02 s^−1^.The molecular brightness was calculated with ImageJ with and without our algorithm detrendr (fully integrated in ImageJ) and a macro to calculate the Brightness (defined as the variance divided by the mean intensity pixel by pixel, available on request).

## RESULTS

To evaluate RH detrending, we compared it with existing techniques in the presence of either (i) various degrees of photobleaching or (ii) fluctuating laser power. We used simulated images of 10,000 diffusing fluorescent particles, sampling at a rate appropriate for number and brightness analysis.^6^ With three different brightnesses (*ϵ* = 0.001, 0.01 and 0.1) which encompass the range of brightnesses we have seen with conventional fluorophores with appropriate acquisition settings – we measured the difference between the brightness calculated with these detrending techniques and the known true brightness from the simulation. The techniques compared to RH were moving average (*boxcar*) smoothing with a constant of 10 frames in either direction, exponential smoothing with automatically chosen parameter^5^ (*autotau*) and no detrend at all (*none*).

Surprisingly, in the presence of photobleaching of constant rate with bleaching by up to 30%, the best choice was no detrend at all. The best of the rest was RH (Figure 1), having errors only slightly worse than *none* and consistently less than half of those of the next best technique, *autotau*. The worst was *boxcar*, with errors always at least three times those of *autotau*.

**Figure 1.**
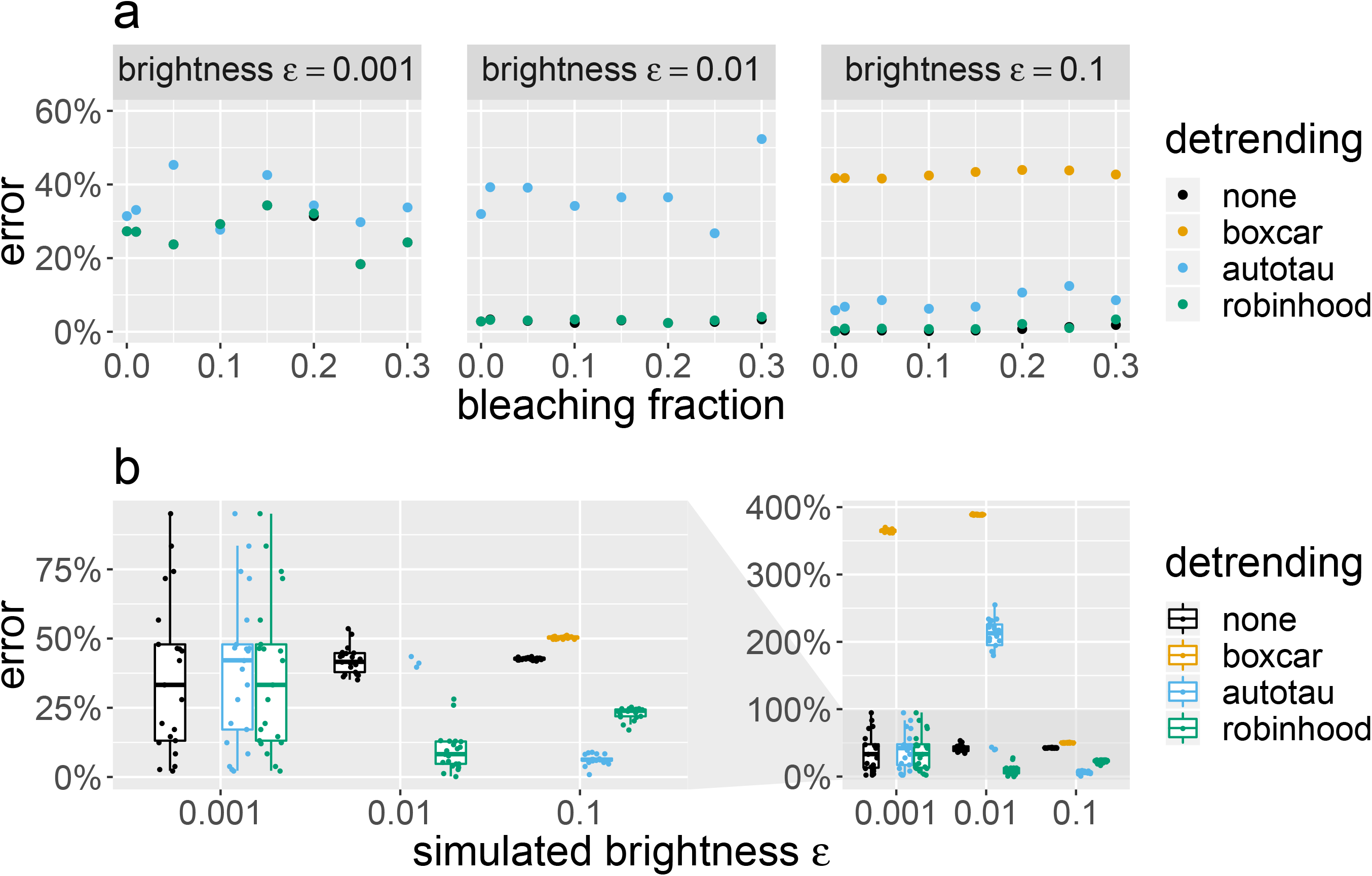
Evaluation of RH algorithm for different brightness as compared to other detrending methods. **a.** Mean relative error in mean brightness ϵ calculation made with different detrending algorithms for different brightnesses and bleaching fractions with constant bleaching rate. For ϵ = 0.001, 0.01, *boxcar* does not appear because its mean relative error is so high (> 200%). For ϵ = 0.001, all of the errors are high because the data contains such little information (only 1 out of every 400 pixels is nonzero), so even with no bleaching and no detrend, there are high statistical errors (> 20%). In all cases, *none* is best, RH is next best and RH almost matches the performance of *none*; all other competitors’ performances are significantly worse. **b.** Mean relative error in brightness ϵ calculation in the presence of a laser which has mean power P but with power fluctuating sinusoidally between 0.5P and 1.5P, measured for different brightnesses ϵ (where ϵ is the brightness at laser power P). Here, RH outperforms *none* in all cases, while for the common low brightness ϵ = 0.01, both *boxcar* and *autotau* have extremely large errors of > 200%. Interestingly, for the high brightness ϵ = 0.1, *autotau* is best and RH is a respectable second.

**Figure 2.**
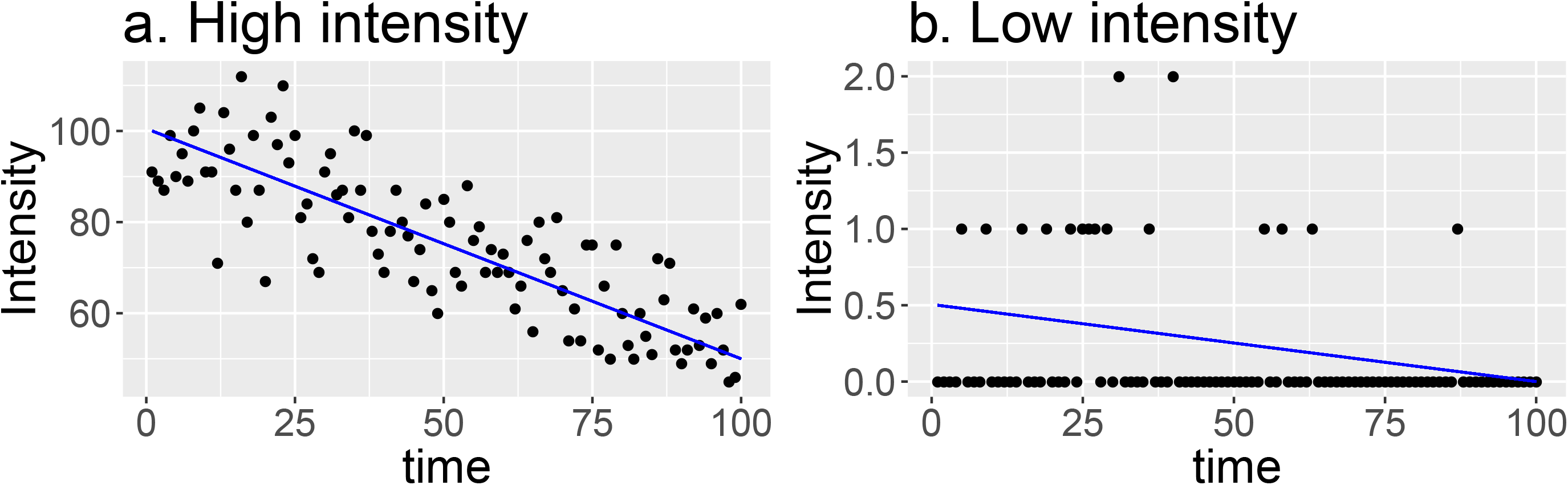
Comparison of two intensity profiles coming from COS-7 cells expressing CCR5-mTFP1. COS-7 expressing CCR5-mTFP1 with a high concentration (left panel) present a clear bleaching profile (blue line). Other COS-7 cells expressing low amounts of CCR5 present many pixel with 0 photon counts and here only RH is able to account for the real photobleaching profile.

In the presence of a laser which has mean power *P* (resulting in fluorophore brightness *ϵ*) but where the power fluctuates sinusoidally between 0.5*P* and 1.5*P* (resulting in fluorophore brightnesses of 0.5*ϵ* and 1.5*ϵ*), the best method was RH (Figure 1), being the only method with errors consistently below 25%. The next best method was *none*, being the only other method with errors consistently below 50%. Notice that at the higher brightness of *ϵ* = 0.1, *autotau* has the best performance, but at the (more common) intermediate brightness of *ϵ* = 0.01, *autotau* has terrible performance, with errors of over 200%. The fact that autotau is best at the high brightness (and hence high intensity) level *ϵ* = 0.1 is noteworthy as it confirms our earlier assertion that smoothing methods (like *autotau*) struggle at low intensities (Figure 3) but not at higher intensities.

In the community, it has long been thought (including by ourselves) that detrending is necessary and that its primary purpose was to correct for bleaching. These simulations show that in the presence of steady bleaching alone, not detrending at all is a fine strategy. We also found that in the presence of sinusoidal variation in mean intensity, not detrending at all has better performance than all currently available detrending methods except for *autotau* (which is best at higher intensities/brightnesses). Hence, we find little evidence in favour of currently available detrending methods. However, we have also shown that in the presence of sinusoidal variation (which cannot come from bleaching), it is better to use Robin Hood detrending than to not detrend at all in all cases. We have also tested the algorithm in live cells expressing CCR5-mTFP1 in the presence and absence of CCL5 (Figure 3). The addition of CCL5 causes a shift in the average brightness from 1.012 to 1.024 (true brightness from 0.12 until 0.24 which is exactly the double and accounts for dimerization). When RH is not employed this dimerization calculation is less precise as the true brightness changes from 0.15 until 0.21.

**Figure. 3.**
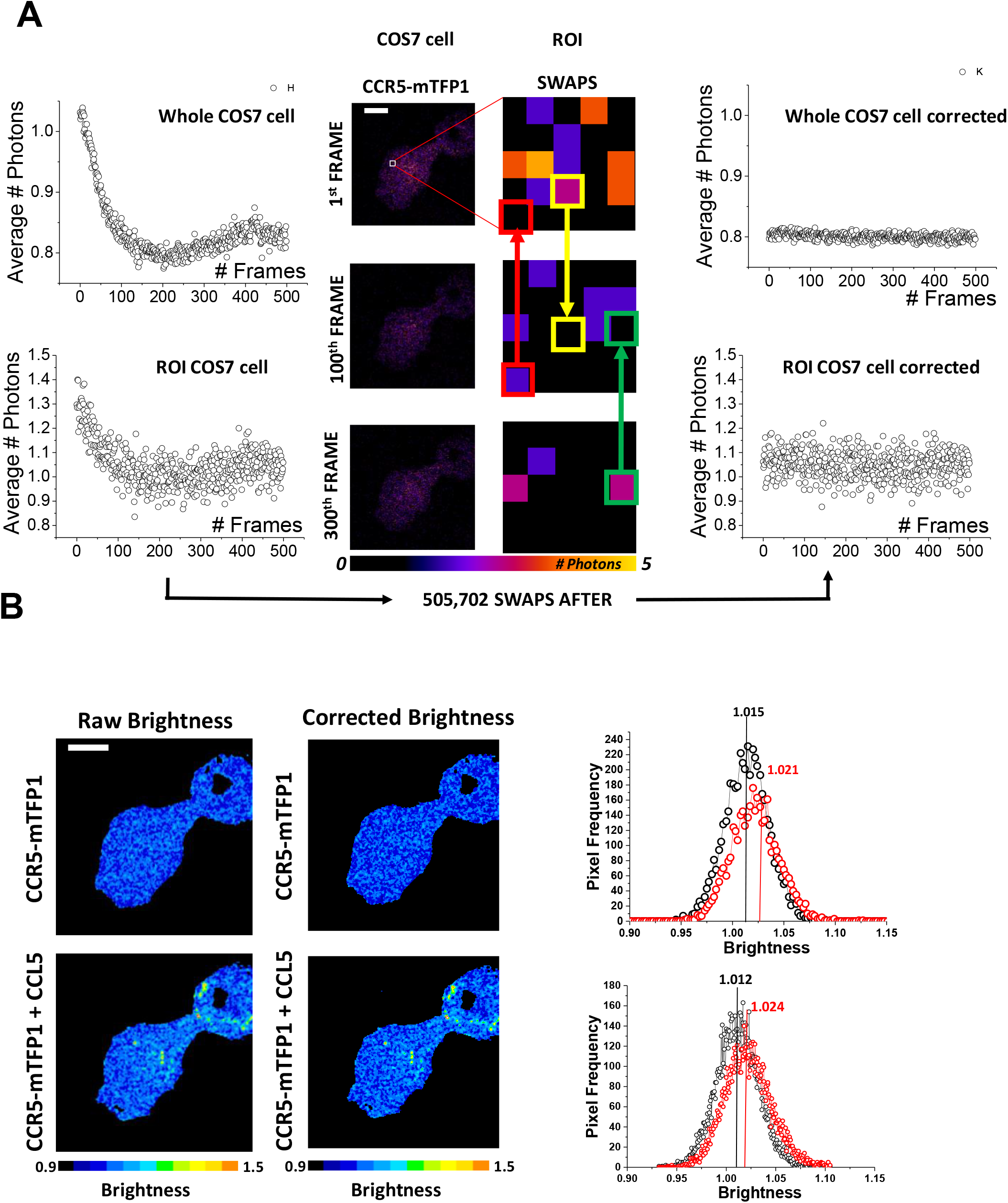
Evaluation of the RH algorithm in live cells. (A) Examples of intensity traces as a function of time coming from regions of interest of COS-7 cells expressing CCR5-mTFP1 before (left panels) and after RH correction (right panels). RH is able to correct both, the average bleached profile coming from all pixels and the small region of interest with very low intensities. (B) COS-7 cell expressing CCR5-mTFP1 before and after addition of ligand CCL5.

In short, we recommend low laser power to reduce photobleaching which leads to lower intensity images. For these low intensity images, we recommend RH detrending. If users find themselves with higher intensity (mean photons per pixel ≥ 10) images, we recommend *autotau* as it will perform better and faster in this case.

## SUPPORTING MATERIAL

This algorithm is available as an ImageJ^7^ plugin (see https://github.com/rorynolan/ij_detrendr#readme) and as an R^8^ package (see https://github.com/rorynolan/detrendr#readme).

## AUTHOR CONTRIBUTIONS

R.N produced the code, run the simulations and the analysis. M.I. provided sample preparation and microscopy acquistions. C.S and Y.J offered funding support. S. P-P. conceived and supervised the project. S.P-P and R.N wrote the manuscript.

## ACKNOWLEDGEMENTS

S.P-P acknowledges funding from the Nuffield Department of Medicine Leadership Fellowship and Gobierno de Espana, Programa Estatal de I+d+I Orientada a los Retos de la Sociedad RTI2018-098415-A-I00. C.S. acknowledges funding from European Research Council grant (647278) and from Cancer Research UK (C20724/A26752). E.Y.J.acknowledges funding from Cancer Research UK and the UK MRC (C375/A17721 and MR/M000141/1) and all authors from the Wellcome Trust Core Award (203141)

